# Unusual traits shape the architecture of the Ig ancestor molecule

**DOI:** 10.1101/2024.07.22.604567

**Authors:** Alejandro Urdiciain, Thomas Madej, Jiyao Wang, James Song, Elena Erausquin, Philippe Youkharibache, Jacinto López-Sagaseta

## Abstract

Understanding the ancestral Ig domain’s molecular structure and tracing the evolution of Ig-like proteins are fundamental components missing from our comprehension of their evolutionary trajectory and function. We have elucidated the molecular structures of two Ig-like proteins from the evolutionary most ancestral *phylum, Porifera*, revealing previously unidentified Ig-domain features that highlight the concomitant presence of a novel Ig “Early Variable” (EV)-set in tandem with a C1-set domain. The latter, to our knowledge, has never been reported before in non-vertebrates. The IgV and IgC1 sets and their combination into functional Ig-like receptors are part of the adaptive immune system in higher vertebrates, which allows for highly specific immune responses. These observations in the ancient *Porifera phylum* could indicate the presence of primitive forms of adaptive immunity or a foundational immune strategy that has been conserved through evolution. By unveiling important clues into the molecular configuration of ancestral Ig domains, these findings challenge and expand our understanding of how immunity has evolved within its current landscape.

## Results and discussion

The sequence homology of the *Geodia Cydonium* sponge Ig-like proteins to antibody domains was noticed early on1–3, but the molecular structure remained unknown. A search for homologous structures using either VAST4,5 or DALI6 led to thousands of structural homologs to the N-terminus domain. This is expected since there are over 13,000 structures in the PDB (protein data bank) containing over ca. 75,000 Ig-domains belonging to the four types of variants: V-set, I-set, C1-set, or C2-set. The overwhelming majority of which consist of antibody fragments: Fabs with IgV-IgC1 of heavy and light chains, or simply VH-VL domains. We therefore looked not just for single domain level structural homologs, but for pairs of Ig-like domains in tandem, and their consecutive domain topologies, in addition to sequence and structure homology, to determine the most relevant evolutionary related two Ig-like domains in tandem among known proteins in the PDB.

Ig-containing regions of both SAML and RTK from *Geodia Cydonium*, found in their extracellular space, were successfully expressed in sf9 cells and purified to homogeneity, as indicated by the elution profiles under size exclusion chromatography (Supplementary Figure 1).

Both SAML and RTK proteins led to thin plate-looking crystals that enabled structure determination with 1.6 and 2.1 Å resolutions, respectively. Overall, both datasets allowed the calculation of electron density maps of enough quality to accurately build the molecular morphologies of both receptors (Fig. 1a). SAML and RTK molecular structures are constituted by elongated assemblies of two Ig-like domains in tandem connected to one another by short and conserved Leu-Pro-Leu linkers. This configuration resembles protein domain architectures common to many cell adhesion and cell surface receptors in vertebrates.

**Figure 1.**
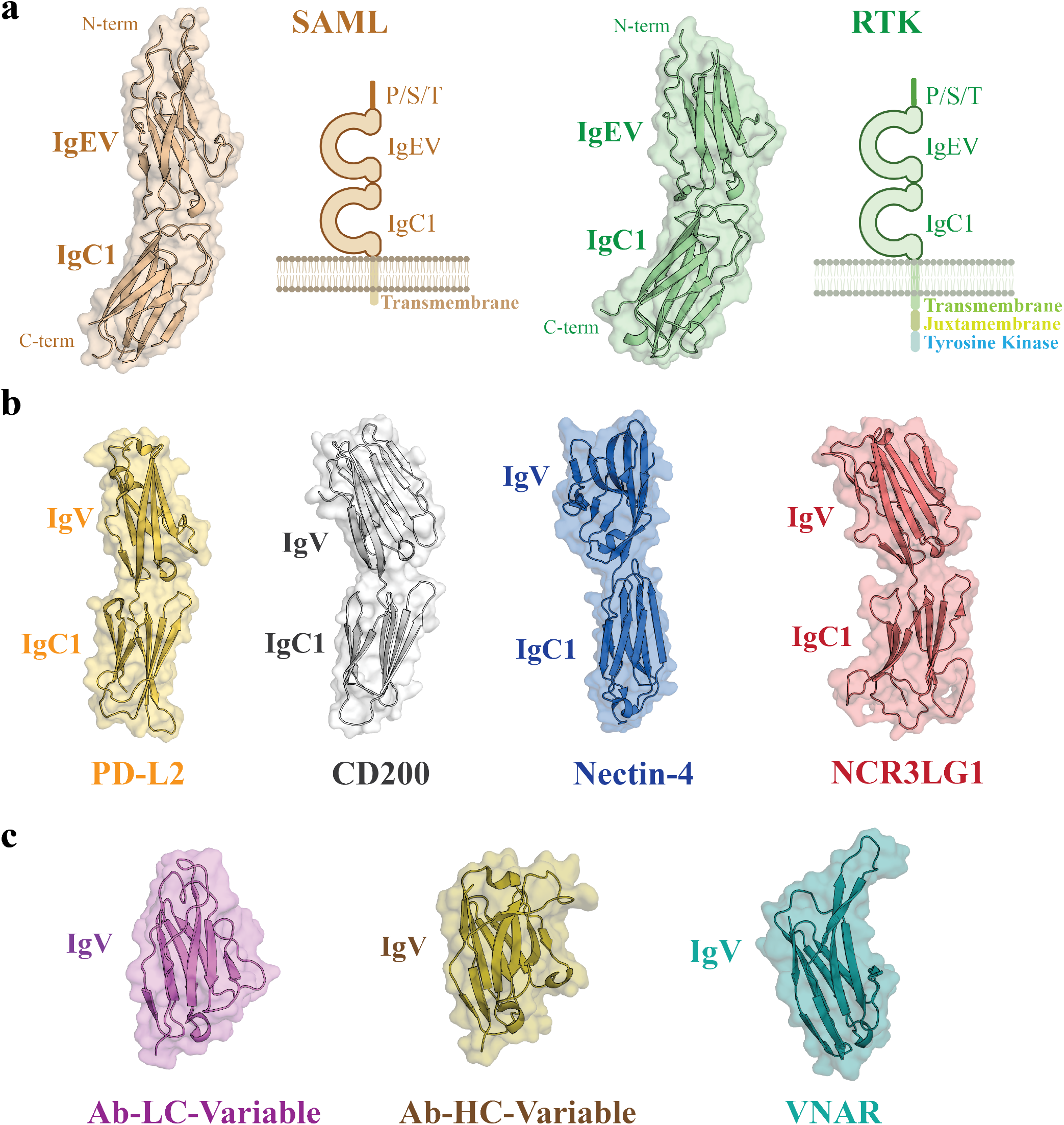
Structure and Ig configuration of SAML and RTK receptors. **a**, the molecular structures of SAML and RTK extracellular Ig-like regions are shown with wheat and pale green colors, respectively, in cartoon and surface mode along with a schematic representation of the corresponding full-length receptors anchored in the cell membrane. N- and C-terminal sites are indicated as well as the transmembrane and intracellular regions, which are assigned accordingly. **b**, display of analogous vertebrate Ig-like receptors configured by an IgV-IgC1 tandem: *PD-L2*, Programmed Cell Death 1 Ligand 2 (pale-yellow color), *CD200*, B-lymphocyte antigen (light gray color), *Nectin4*, Nectin cell adhesion molecule-4 (blue color) and *-B7-H6(NCR3LG1)*, a ligand to Natural cytotoxicity triggering receptor 3 also known as NKp30 or CD337 (salmon color). **c**, the structures of three different Ig-V sets are also shown for structural comparison purposes: *Ab-LC-variable*, antibody light chain variable region (violet color), *Ab-HC-variable*, antibody heavy chain variable region (olive color), *VNAR*, shark variable domain of new antigen receptor (teal color). Pro-Ser-Thr(P/S/T)-rich domain.

Interestingly, some of the structural homologs with the highest sequence identity up to 30% point to human IgVL lambda (PDB 3T0X), IgVH (PDB 3N9G) domains, and 20% with Shark VNAR (Variable domain of New Antigen Receptor) domains (PDB 1VES) (Fig. 1b). Also, VAST structural homologs encompassing N- and C-terminus Ig-domains in tandem with good overall (rigid) structural alignments, lead to PD-L2 (PDB 3BP5), CD200 (PDB 4BFI), Nectin-4 (PDB 4FRW), and NCR3LG1 (B7-H6) (PDB 6YJP) that all present an IgV-IgC1 extracellular chain architecture, characteristic of many vertebrate immune cell surface receptors (Fig. 1c). Precisely, the extracellular architecture is composed of two Ig-domains in tandem, the N-terminus with a topology that can be classified, preliminarily, as a V-set followed by a C-terminus domain belonging to the C1-set (Fig. 1a). The orientation of the Ig domains of SAML/RTK depicts a single polypeptide conformed by two β−sandwich barrel-like folds subtly angled in relation to each other.

Each of the two Ig-domains in SAML and RTK present the Cys-Cys-Trp canonical triad, highly conserved across the vast array of Ig-domains7–10. Thus, the morphology of these Ig-like domains presents two confronted β-pleated sheets held together by a disulfide bridge and a neighboring Trp residue that contributes to the overall compactness and stability of the Ig fold (Supplementary Figure 2).

An additional Trp is located in the N-terminal domains of SAML and RTK, near the disulfide bridge, presumably to further balance the overall architecture of these domains. However, this Trp is not conserved in other Ig-like proteins in vertebrates, which might indicate a less relevant structural role and subsequent loss through evolution.

The combination of a monomeric IgV-IgC1 tandem extracellular region points to an ancestral form equivalent to the immune cell surface receptors of the vertebrate immune system. There are naturally both similarities and differences between the vertebrate V-set domains found in antibodies and the SAML/RTK N-terminal domains, which resemble more to a Shark VNAR domain, but also to some human V-set domains as found in PD-L2.

### The N-terminal domain

The N-terminal domain topology can be compared to Ig-domains of either V-set or I-set. Indeed, many of the immunoglobulin superfamily domains in cell adhesion molecules and surface receptors belong to the I-set type11. Yet, most cell surface receptors in vertebrate immune cells tend to have an IgV domain designed to interact with their ligands IgV domains, as for receptors-ligands such as PD-1, PD-L1, PD-L2, CD28 and CTLA-4 of the B7:CD28 extended family, that contain one or two extracellular Ig-domains with an IgV-domain at the N-terminus. V-set and I-set topologies are extremely close when considering the Ig-framework alone11. Importantly, the most significant structural differences between the two types lies in the loops that are very short in I-set domains as compared to their V-set counterparts (Fig. 2). These differences are particularly notorious for the short FG loop, which correspond to CDR3 in antibodies, the BC loop, CDR1 in antibodies, and a C’D loop with the absence of a C” strand and consequently no C’C”, CDR2 in antibodies (Fig. 2 and Supplementary Figure 3). Importantly, the FG loop in SAML/RTK represents a manifest divergence as to potentially exclude SAML/RTK N-terminal domain from belonging to the Ig-I set. The FG loop length in SAML/RTK resembles that of VNAR, falling in between the length range observed for antibody light and heavy chains CDR3 loops (Fig. 2). Thus, the relatively extended length of SAML/RTK CDR3-like loop is especially interesting in pointing to a potential functional role already present in primitive organisms.

**Figure 2.**
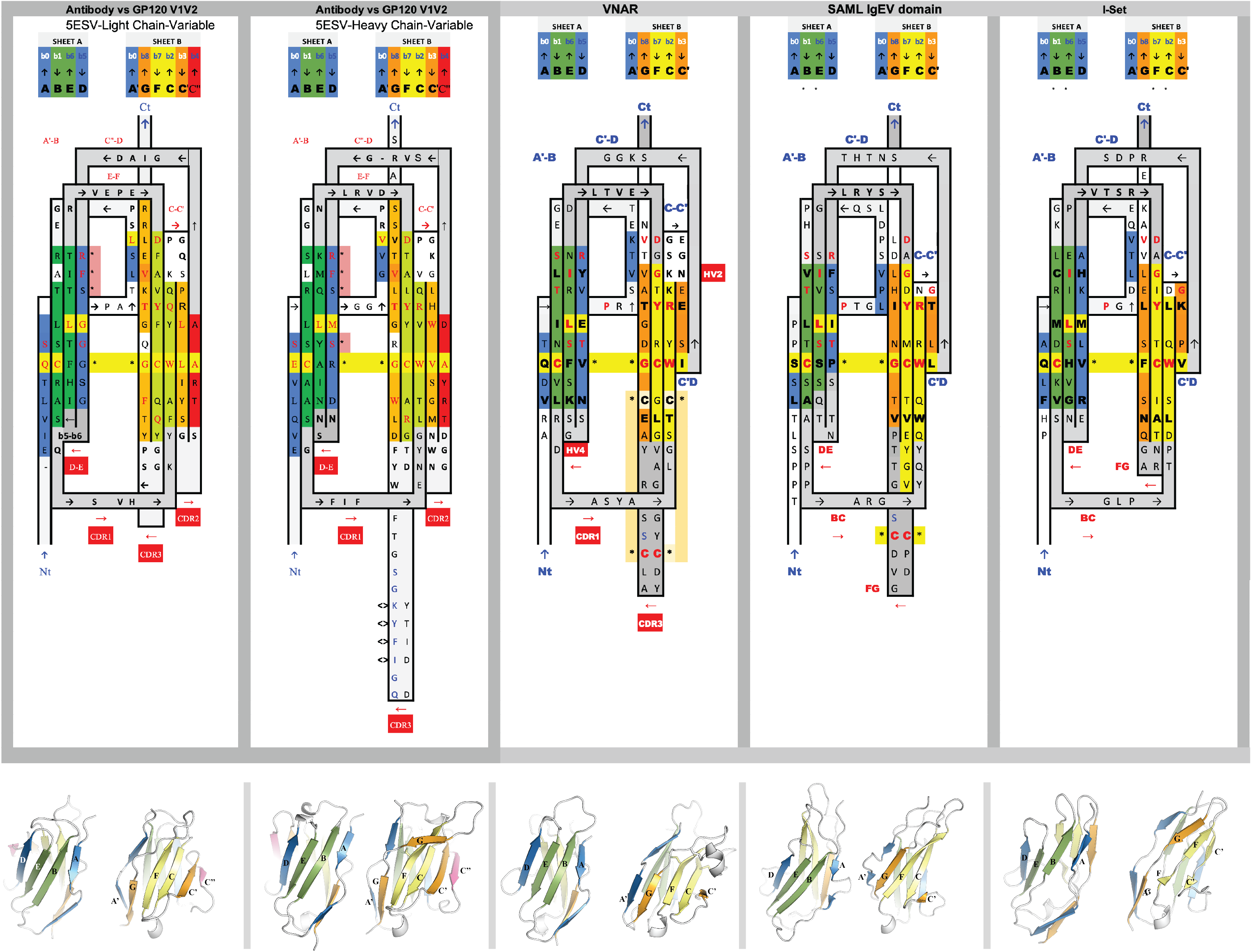
Sequence and topology 2D alignment of N-term Ig-domains. 2D sequence/topology alignments allow integration of sequence and topological assembly into one single map and subsequent comparison of a series of diverse Ig-sets. Shown are the 2D maps for a series of Ig-sets representative of the evolution the Ig-domain in the animal kingdom. Colors correspond to the *Ig Protodomain reduced rainbow color spectrum*^14^ with four strands colors blue, green, yellow, orange for ABCC’ and DEFG and a fith color red for the C” strand, if present. Using this color scheme, a sheet ABED will be blue-green-green-blue; a sheet GFCC’ will be orange-yellow-yellow-orange.

One should note however that IgV domains in sharks, called VNAR, lack the CDR2 region too, called in that case HV2, still hypervariable in sequence. VNAR domains12 and IgI domains align in fact remarkably well as noticed previously13. This region can also be either highly flexible in well-known human cell surface receptors, such as PD-L1, or missing altogether, as in the case of PD-L2. As seen in Figure 2, the N-term domains of SAML/RTK show significant sequence similarities with both IgI and VNAR domains.

The three share a significant analogy in their respective topologies, while all differences between IgI and VNAR, as with SAML/RTK, occur in loops’ length and structure. In conclusion, SAML and RTK N-terminal domains resemble VNAR and PD-L2 in that respect. SAML/RTK show significant similarities on central strands B (SxTxxC), C (WxR) on the ABED sheet, E including the EF loop (SxSLxISxxLRxxx) and F (DxGxYxC) on the GFCC’ sheet to both. While both VNAR and IgI domains exhibit the highly conserved pattern in strand F, the conservation on the preceding E strand and EF loop is more extensive and convincing with VNARs than with IgI. An additional sequence homology with VNAR and antibody variable domains also occurs in the D-strand (RF/YxxT/S) (Fig. 2).

The presence of two cysteines structurally configured to form a disulfide bridge in the CDR3 loops occurs in natural antibodies’ VH domains14. Non-canonical cysteines are also common CDR3 loops of VNAR type I-III domains15. The only loop where SAML/RTK resemble an IgI is the CC’ loop, that is 4-aa longer in VNAR. However, this discrepancy with VNAR suggests that SAML/RTK Ig domains should not be considered as of pure VNAR type.

In addition to these observations, SAML/RTK N-terminal domains also present an AA’ strand break, with A being part of the ABED sheet and A’ part of the A’GFCC’ sheet, as occurs in both I- and V-sets. Thus, unlike the canonical IgV-set, SAML/RTK N-terminal Ig-domains do not feature a C″ strand. The absence of such strand is a known feature in the IgI-set, and therefore this feature imprints a partial Ig-I set character to SAML/RTK Ig N-terminal domains. Further, in both V-set and I-set domains, the Ig domain begins with an A strand in the ABED sheet and transitions through a structural kink into the A′ strand. This A′ strand interfaces with the GFCC′ sheet and frequently exhibits a kink associated with a cis-Proline residue, running in parallel with the G strand to protect it from exposure. However, the AA’ strand split in SAML/RTK exhibits a remarkable bulge than what is observed for IgI or IgV domains (Supplementary Figure 3). This unique feature suggests that the AA’ strand associates with functional properties (Richardson and Richardson 2002; Benian and Mayans 2015) that might have been lost through evolution, and might well constitute a further distinctive trait associated with the ancient *Porifera* Ig domain. Also, the A’B-loop in SAML/RTK is more extended than in IgV and IgI domains alike. In SAML, we find the A’B loop is conformed by a 4-aa (LSQP) motif, while only 2-aa patterns are detected in VNAR and the I-set, TG and EG, respectively. Additionally, we also noted that the N-terminal regions of SAML/RTK feature a proline-rich A strand, a trait commonly observed in I-set and V-set domains. On the other hand, however, these regions lack key I-set domain hallmarks: specifically, the motif where three residues follow the YxC motif in the F strand, with an Asn residue serving as a connector between the BC and FG loops17 (Fig. 2). Thus, both the AA’ protrusion and extension in the A’B loop might represent unique features of ancestral Ig-like domains. Based on our observations, this domain is not an IgI from a sequence or loop structure standpoint.

Accordingly, and if considered an Ig-V similar to VNAR, it might well represent a novel domain that we accordingly name “*Early Variable” (EV) domain”*, with the major difference to most IgV domains being larger AA’ protrusion, and a longer A’B loop. In addition, a short C’ accompanied by a lack of C” strand and CDR2 region, a pattern shared with VNAR domains, excludes SAML/RTK N-terminal domains from belonging to the canonical Ig-V domain family. Combined with a long CDR3 (FG) loop and the absence of a CC’ loop, these findings further strengthen the concept of a novel and ancestral Ig-domain.

A comparison of those regions equivalent to the CDR loops in antibodies and TCRs show a remarkable degree of structural conservation and overall positioning throughout primitive and contemporary organisms (Supplementary Figure 4). This highlights a fine-tuned evolution of the Ig domain defined by a strong preservation of the Ig beta-sandwich architecture scaffold accompanied by structural transitions involving not only molecular diversification but the emergence of novel mechanisms, i.e., somatic hypermutation.

### The C-terminal domain

The C-terminal domain however presents a clear C1-set topology and this is a significant finding. Effectively, while V-set domains have been found in numerous life forms, the C1-set has only been found in vertebrates so far18 and is considered a hallmark of adaptive evolution in vertebrates10. This represents a surprise since Ig-C1 domains have been linked to the evolution of the vertebrate immune system18 and is present in particular in molecules of the adaptative immune system (antibodies, TCRs and MHCs). It is found in numerous cell surface receptors of the innate immune system such as B7 proteins such as PD-L1/L2 or CD80/86, and present especially in BCRs, antibodies, TCRs and MHC molecules involved in adaptive immunity jawed vertebrates19–21 (Fig. 1) Thus, the C1-set domains were believed to be present only in vertebrates.

Importantly, in addition to variable domains, the Ig-C1 architecture confers antibody chains the ability to dimerize. This is believed to be a one of the innovations in the adaptive immune system10. Effectively, C1-set domains are found to dimerize in an antiparallel mode, using the ABED sheet in antibodies’CH1/CL domains, as well as CH3 or CH4 domains in polymeric Igs22,23, or in TCRα/β. In shark antibodies (IgNAR), the C1-set constant domains that follow the VNAR variable domain also dimerize through their C1 and C3 domains12,24. Neither SAML nor RTK C-domains have shown evidence for dimerization in either their crystal forms or in solution, as confirmed by crystallographic and size exclusion chromatography with multi-angle static light scattering (SEC-MALS) studies (Supplementary Figure 5).

Therefore, the presence of the C1-set domain, apart from establishing an evolutionary link to vertebrate domains, may point to a possible oligomerization *in vivo*. Alternatively, it might indicate an additional evolutionary step that confers the C1-set the potential to dimerize in the more evolved forms of life.

Immunoglobulin-like (Ig) domains are classified into four primary types: IgI, IgV, IgC1, and IgC2, each uniquely characterized by the presence or absence of additional β strands17,25–27. The structural configuration and β strand composition distinguish IgV and IgC domains. IgV domains are typically composed of nine β strands, organized into two β sheets: the first sheet includes strands A, B, E, and D, while the second comprises strands G, F, C, C’, and C” (the region covering C” is sometimes replaced by a flexible loop as in PD-1, or missing as in PD-L2, or VNAR domains found in Sharks. The A strand can be split into two segments, that are conventionally named A and A’, for their association with either side of the sandwich sheets. In contrast, IgC-set domains, which are subdivided into IgC1 and IgC2, feature a more compact structure with fewer β strands. The IgC1 variant omits the C’ and C” strands, presenting only seven β strands. IgC2 domains incorporate an extra C’ strand, akin to that in the IgV domains but lack the D strand found in the back face of IgV structures. This difference alters the configuration and surface area of the β sheets between the IgC1 and IgC2 types, despite their similar strand count.

Thus, upon analysis of the canonical four main Ig sets, it is determined that SAML/RTK C-terminal domains belong to the IgC1-set, distinguished by the inclusion of a D strand and the absence of a C’ strand (Figure 3). The SAML/RTK domain features specific motifs: LxCxA on the B strand, NxxxSLxI on the E strand, CxxxS on the C strand, and YxCxV on the F strand, aligning with the hallmark characteristics of canonical IgC1-set domains.

**Figure 3.**
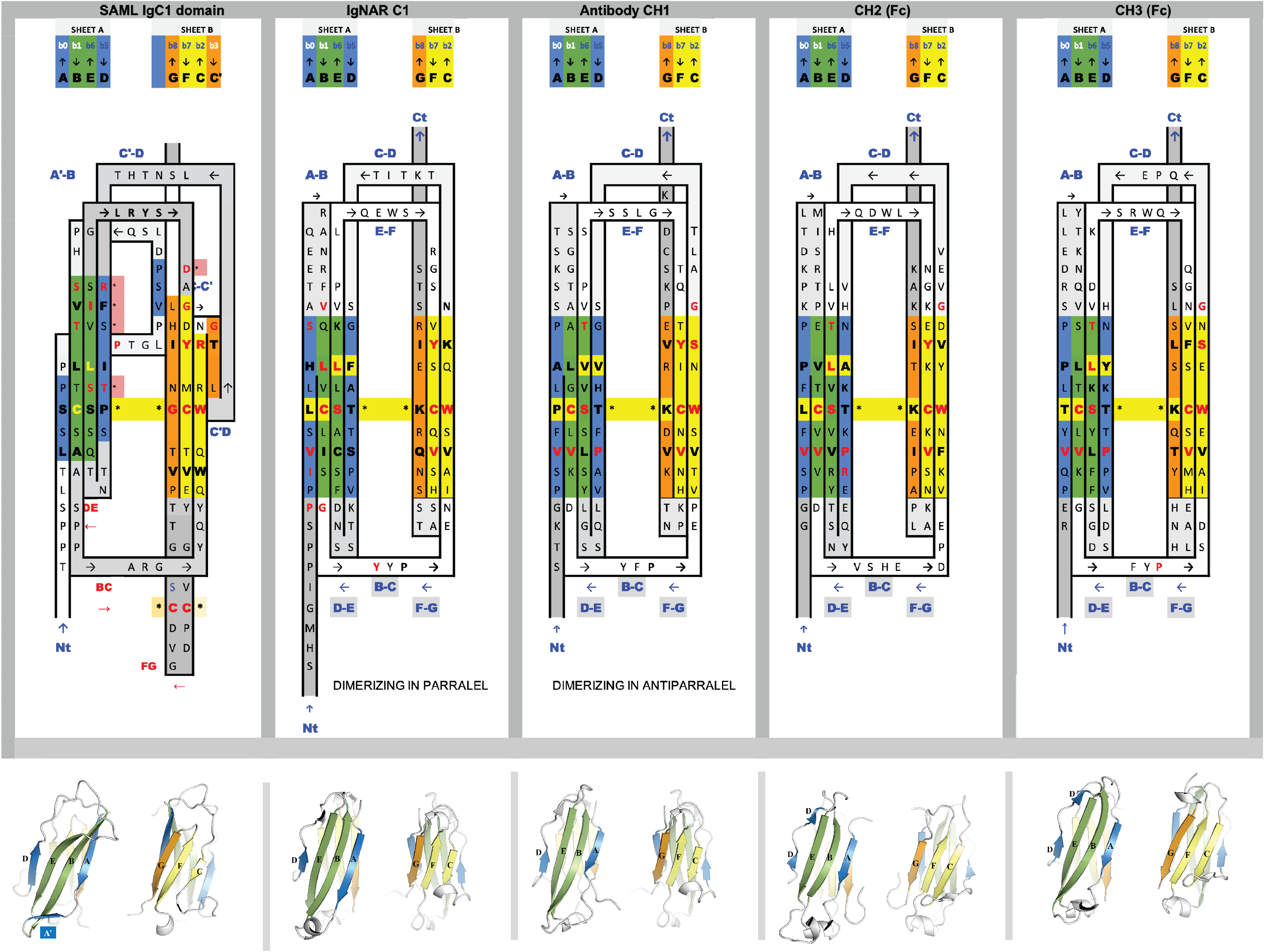
Sequence and topology 2D alignment of N-term Ig-domains. 2D sequence/topology alignments allow integration of sequence and topological assembly into one single map and subsequent comparison of a series of diverse Ig-sets. Shown are the 2D maps for a series of Ig-sets representative of the evolution the Ig-domain in the animal kingdom.

Interestingly, these findings align with the hypothesis by Pasquier and Chrétien28 pointing to a primitive receptor form of monomeric nature, assembled by Ig-domains without somatic recombination and featuring adhesion molecule-like properties28.

A distinctive feature of the interdomain interface of SAML and RTK is the C1 domain presenting a long DE loop forming a compact supersecondary structure with the BC loop, something quite unusual in known IgC1 domains, indeed looking quite similar to a C’E loop as observed in C2-domains straddling the two sheets of the sandwich, while stemming from the D strand. An IgC1 vs. IgC2 topology lies essentially in swapping the C’ strand in IgC2 to a D strand position hence forming an IgC1 sandwich composed of GFC and ABED sheets rather than GFCC’ and ABE sheets in IgC2. The prominent compact BC and DE loop of the C-terminal Ig domain (IgC1) domain form a compact substructure, that in turn forms a compact interface with the A’B and EF loops of the N-terminal Ig domain (IgEV), with a buried surface of ca. 500 A^2^. This does confer to the IgEV-IgC1 tandem domains of SAML and RTK a fused/rigid structure.

### Ig-tandemers

Rigid structural alignments can find the structural homologs of two domain structures only if their relative domain orientation is similar, and protein chains containing multiple Ig-domains exhibit a very high conformational plasticity, due to flexible interdomain linkers and diverse loops involved at the domain-domain interfaces. To identify the closest homologs among known structures, we looked for structures with two consecutive Ig-like domains in tandem, considering a rigid body alignment at the domain level, independently of their relative orientation. We call such structures Ig-tandemers. In our case, we can focus on those tandemers presenting a C-terminus domain with a C1-set topology Ig while N-terminus domain may present either an I-set or V-set topology. A canonical C1 topology presents a straight A strand and the presence of a D-strand as a distinctive mark.

We used the NCBI structure alignment program VAST4,29 to compare the SAML structure against the full PDB database (as of Jan 31, 2024). The two SAML domains were run separately as also the full chain. We then combined the results by extracting those protein chains that aligned with the two SAML domains, in tandem. For each domain we computed the alignment length, RMSD for that structural superposition, and the number of identical residues in the SAML/target alignment. We ended up with 18047 hits from 6581 different PDB files (The full list is available as supplementary data). The chained Ig-domains across the PDB structures offer a diversity of tandemer topologies presenting variants from IgV-IgC1, IgV-IgC2, IgI-IgI, IgI-IgC2, etc, with diverse relative orientations (not considered explicitly).

Given the overrepresentation of antibody Fab and TCR structures in the PDB database that possess an IgV-IgC1 topology, we separated all the hits into a Fab antibody/TCR dataset and a non-antibody dataset. The former containing ca. 15,000 chains and the second with ca. 3,000 chains. Sequence homology to represented Fab antibodies/TCRs averages 23.8% (max 47.7% on structurally aligned residues) for the N-terminal domain hits (Ig1) and 15.8% on the C-terminal domain (Ig2) (max 32.8%). Moreover, for the Ig1 domain an average of 76 residues were aligned with a 2.1A average RMSD, and for the Ig2 domain an average of 63 residues with a 2.0 Å average RMSD. In the the non-Ab dataset the sequence identity averages 16.4% (max 35.2%) on Ig1 and 16.6% (max 34.4%) on Ig2. For Ig1 there was an average of 71 residues aligned and a 2.3 Å average RMSD, for Ig2 an average of 59 residues aligned and 2.1 Å average RMSD. In this dataset, considering sequence, topology and structure altogether, VAST identifies several tandemer homologs, pointing to type I cell surface receptors involved in cell-cell adhesion and immunity.

These proteins are all type I cell surface receptors, as are SAML and RTK. Among them CD200, PD-L2 (B7-DC, CD273), NCR3LG1(B7-H6), CD80 (B7-1) and nectin-4. CD200 (4BFI.B) is expressed on a wide range of cell types, including neurons, endothelial cells, and immune cells. It interacts with its receptor, CD200R (4BFI.A); PD-L2 (3BP5.B) is expressed on activated human T cells and regulates their function30 and B7-H6 (P.C) is also part of the B7 family31 that consists of structurally related cell-surface protein ligands, which bind to receptors on lymphocytes that regulate immune responses. CD80 (B7.1) (1I8L.A) binds to CD28 and CTLA-4 receptors on T cells in forming and stabilizing the immunological synapse. All these interacting cell surface proteins are part of an extended B7:CD28 family32. Nectins are adherens junction proteins involved in the nervous and the immune system, and also serve as virus receptors, such as the well-known poliovirus receptor (PVR or CD155), or nectin-4 (4FRW.B) as the measles virus receptor33. We show a compendium of more structural homologs in Supplementary Table 1, such as the Lymphocyte activation gene 3 LAG-3 (7TZE.A), an immune checkpoint receptor and target for cancer immunotherapy34–36 or CD277/butyrophilin-3 (4F9L.A)37. A more comprehensive list of hits is presented in Supplementary Table 2.

### Disulfide bridge in the “CDR3” loop

Given the quality of the electron density signal, we could notice a remarkable trait in SAML Ig-EV domain. A cysteine pair in the FG loop (“CDR3”), where Cys252 presents up to three alternative conformations, of which one leads to a disulfide bond with Cys258 (Supplementary Figure 7). Occupancies and B-factors for the three conformations were determined, respectively, to values of 0.31/23.94 (conformer 1), 0.25/22.22 (conformer 2) and 0.44/22.13 (conformer 3) through anisotropic refinement of B-factors. Non-canonical cysteines have been observed in human immunoglobulins, particularly in the CDR3 of the heavy chain (CDR-H3), as inferred from a study with ∼3 billion VH sequences38.

The number of non-canonical cysteines, i.e., those found outside the Ig-fold main body and which are encoded by Diversity Gene segments, in the most common human CDR-H3 can vary from 1 to 8 cysteines, and their frequency decreases as the number of cysteines increases. It has also been observed that the size of the CDR-H3 grows along with the number of non-canonical cysteines. The non-canonical disulfide bridge is more common in other species compared to humans, specifically in rooster, camel, llama, shark and cow, and are more frequently observed in the IgM class. A comparison in Ig sequences between naive lymphocytes and memory lymphocytes revealed there is 2% fewer cysteines in CDR-H3 compared to memory lymphocytes, so it is suggested that this increase in cysteines might be linked to somatic hypermutation.

Here we are talking about an adaptive response, since lymphocytes are involved. In the case of sponges, we would be talking about the innate response, since it is the sequence that it has in itself, and could therefore be involved in the recognition of self and foreign antigens. As for humans, it has been observed that a disulfide bridge can also be formed between a cysteine found in the CDR-H3 and another cysteine found in other region of the Ig, as in CDR-H1 and H2^39^. In a study by Almagro and Colleagues^15^, the authors observed that antigen binding was diminished in a significant manner when the non-canonical CDR-H3 cysteines were mutated to alanine, which indicates the potential relevance of these class of disulfide bridges in antigen recognition. The most common sequence in which these non-canonical cysteines occur in CDR-H3 is: C-4X-C, with 4X being a 4 amino acid sequence, the most prevalent sequences being SSTS or SGGS. In the case of RTK and SAMS, the sequence is C-5X-C (CPDGVDC).

Overall, the observation of multiple conformations for a cysteine residue, with one such conformations forming a disulfide bridge with a neighboring cysteine, highlights the dynamic nature of the FG loop. We hypothesize a structural mechanism that provides this loop with capability to undergo conformational changes that would ultimately confer these ancestral receptors a broader scope with regard to antigen recognition diversity.

### Polymorphism

An important feature previously described for RTK is its polymorphic nature^2^. Polymorphism in molecules of the immune system lead to variations in their structure and functions among individuals, affecting their immune responses. One of the most well-known examples of polymorphism is within the major histocompatibility complex (MHC) genes, found in more evolved forms of life such as vertebrates, including birds, reptiles, and fish. Polymorphisms within MHC genes result in a diverse array of MHC molecules, allowing individuals to recognize and respond to a wide range of pathogens. The presence of polymorphism in proteins of *Porifera*, the most ancient phylum of the Metazoa phylogenetically, along with the lack of the cellular machinery for somatic recombination and gene diversity points to a possible hallmark of the less evolved primitive organisms. To explore a potential interplay between polymorphism and location in RTK, we located the polymorphic positions in RTK structure (Fig. 4). However, we found that polymorphisms are not confined to particular regions. These are instead widely distributed in RTK, with equal frequencies in both N- and C-terminal Ig domains. Polymorphic sites are equally found in loops and in more rigid areas such as the largely populated β sheet strands.

Thus, no evident clues can be inferred that could attribute a particular region in these Ig domains a leading functional role in terms of antigen recognition or cell-cell adhesion mechanisms. Perhaps, the distribution of polymorphic sites in Ig molecules have evolved from a primitive, broad distribution towards a more antigen-specific and topologically concentrated configuration, as occurs in the hypervariable CDRs of the more evolved antibodies and TCRs Ig proteins. In conclusion, it could be argued that polymorphism in proteins of ancient organisms likely mirrors a dynamic interplay between environmental pressures, genetic variation and evolutionary processes, raising a foundation for resilience and adaptation in primitive forms of life.

**Figure 4.**
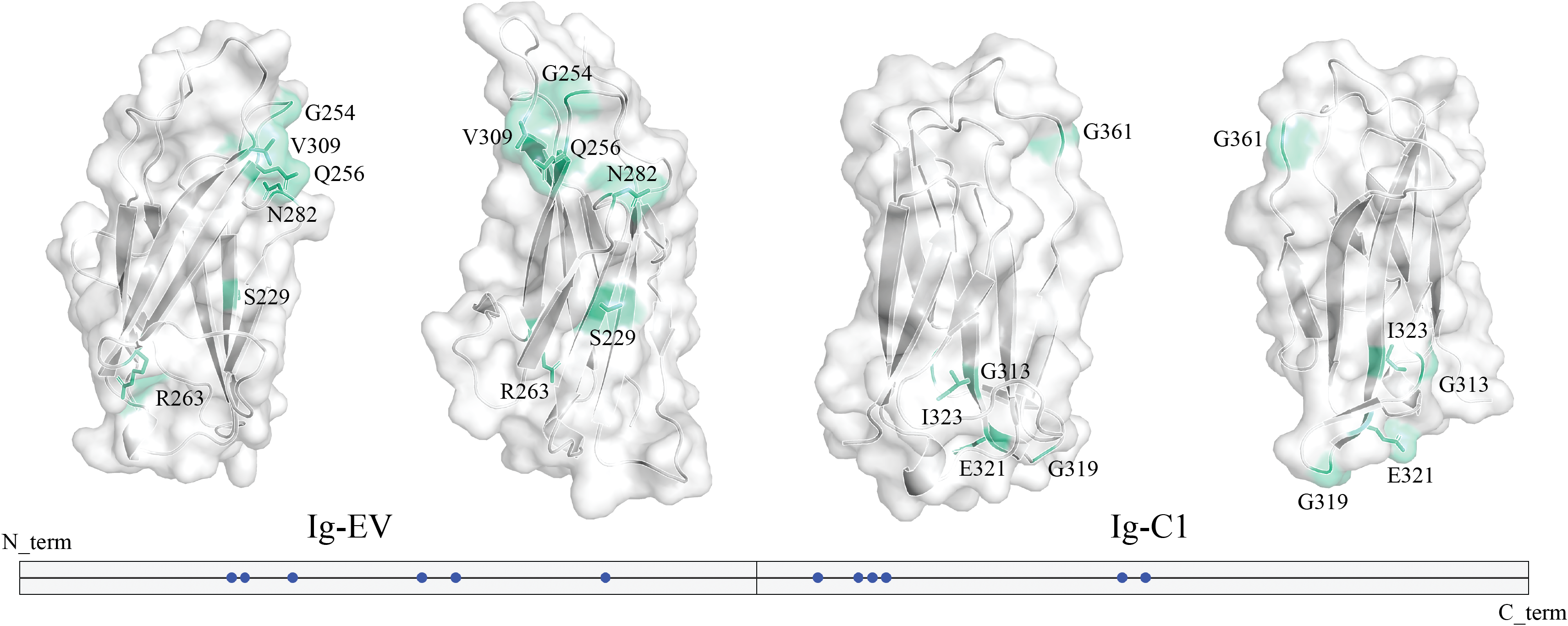
Location of polymorphic sites. A cartoon and surface representation are shown for SAML Ig-EV and RTK Ig-C1 domains, with the polymorphic sites highlighted in greencyan color. The diagram indicates the location of the polymorphic residues (blue circles) along the whole ectodomain of RTK. Those regions that could not be built for RTK due to poor or negligible electron density signal have been replaced by their equivalents in SAML. Numbering of residues is indicated according to the PDB (8OVQ for SAML and 8QPX for RTK).

### Striking loop and PTM (N-glycosylation)

The C’D loop drew our attention due to its length and conformation, folded upon itself and packed against the N-terminal Ig-body (Supplementary Figure 8). The presence of several solvent-exposed and polar residues points to a potential functional role, such as antigen recognition or cell-cell adhesion. Interestingly, three conserved residues, Arg205 and Arg216 in the C’D-loop and Arg236 in the EF loop create a pocket with a net positive charge (Supplementary Figure 9). Another interesting observation in this region is the presence of sugar moiety in the form of N-glycosylation at Asn206. Nevertheless, a heavier N-glycosylation pattern is observed in the C-C’ segment of RTK, with a branched sugar chain common in eukaryotic proteins (Supplementary Figure 10). Altogether, these findings suggest potential roles associated with antigen recognition or cell-cell adhesion mechanisms^40^.

## Methods

### Gene synthesis, cloning and plasmid purification

Codon optimized sequences for sponge RTK (with position 129 being the N-terminal native amino acid) and SAML (with position 157 being the N-terminal native amino acid) containing a N-terminal Twin-Strep tag followed by human rhinovirus 3C protease site were synthesized and inserted in a pUC57-BsaI-Free vector by Gene Universal (Newark, DE, USA). Target sequences were flanked by BamHI and NotI restriction sites. The sponge genes were obtained by digestion with FastDigest Restriction Enzymes (Fisher Scientific) and cloned into the pAcGP67A vector using Optizyme™ T4 DNA Ligase (Thermo Fisher Scientific). The three Ig-like genes were inserted downstream of the gp67 secretion signal sequence included in the pAcGP67A baculovirus transfer vector to facilitate the isolation of the recombinant protein from the supernatant. *E*.*coli* DH5α (Invitrogen) were transformed with the plasmids obtained after the ligation and were incubated overnight at 37 °C. A colony was growth on LB Broth (Lennox) medium overnight at 37 °C in a shaking incubator. Plasmids were obtained using GeneJET Plasmid Miniprep Kit (Thermo Fisher Scientific) following manufacturer’s instruction. Plasmids were purified in order to transduce Sf9 insect cells. Briefly, 1/10 volume of 3M sodium acetate pH 5.2 was added. Then, three volumes of absolute ethanol were added and incubated on ice for 15 min. Samples were centrifuged (14,000g, 30 min at 4°C). Supernatant was discarded, and DNA pellet was washed with 1 ml of 70% ethanol. Samples were centrifuged again for 15 min. Supernatant was discarded and pellet was resuspended on 20 µl of 10 mM Tris pH 8.5. Plasmids were stored at -20 °C.

### Generation of P0 and P1 baculovirus

Sequences were validated by Sanger sequencing. Then, 2×10^6^ Sf9 insect cells (Gibco) were transducted with 500 ng of each transfer plasmid, 100 ng BestBac 2.0 Δ v-cath/chiA Linearized Baculovirus DNA (Chimigen) and 1,2 µl TransIT (Mirus Bio) to produce final recombinant baculovirus. First, recombinant plasmid, BestBac and TransIT were mixed in 100 µl Sf-900 III SFM (ThermoFisher) and incubated for 20 minutes at RT. Once cells were observed to be attached to the well, 1.8 ml of Sf-900 III SFM without FBS nor Penicillin/Streptomycin and the transduction mix was added, and incubated for 5 h at 28 °C. Then, cells were supplemented with 100 µl of FBS and 5 µl of antibiotics. Cells were incubated for 5 days at 28 °C. After that, P0 baculoviruses were collected in the supernatant after centrifugation (300g, 10 min, 4 °C) and stored at 4 °C.

In order to amplify virus titers, new 25 ml Sf9 cultures in Sf-900 III SFM supplemented with 10% FBS and 0.25% Penicillin/Streptomycin, at a density of 1×10^6^ cells/ml were infected with 25 µl of P0 baculovirus. Culture cell morphology and number were observed daily. 48 h after infection, cells were unable to proliferate and its morphology changed, looking swollen. Then, cells were left for another 24 h on an orbital oscillator at 28 °C. Supernatants, containing P1 baculovirus, were collected by centrifugation (3000 rpm, 15 min, 4 °C) and stored at 4 °C.

### Recombinant protein expression and purification

Sf9 insect cells at a density of 2×10^6^ cells/ml in Xpress medium were infected with TwinStrep-tagged recombinant protein P1 baculoviruses at a 1:2000 ratio, and were incubated for 72 h at 28 °C in a shaker incubator. Supernatant-containing protein was collected by centrifugation (10,000g, 15 min, 4 °C), and supplemented with 100 mM HEPES, 150 mM NaCl and 1 mM EDTA, pH 7.4. Recombinant proteins were purified from the supernatant with a StrepTactin 4Flow 5 ml cartridge (Iba Lifesciences) following manufacturer’s buffers and instructions. Elution was concentrated in a 10 kDa 4 ml amicon (Merck) and TwinStrep-tagged protein was digested with 3C protease overnight at 4 °C at a 1:50 w/w ratio. 3C protease-digested protein was purified by size-exclusion chromatography using a Superdex 75 10/300 GL (Cytiva) in Tris-buffered saline (TBS) pH 7.4. Peak belonging to 3C-digested protein was concentrated using an amicon of 10 kDa and 4 ml. 3C protease, which contained a 6xHis tag, was removed from the sample with Nickel NTA agarose resin (ABT). Purified Ig-like proteins were concentrated using 10 kDa Nanosep columns (Pall Corporation), aliquoted and frozen in liquid nitrogen prior to storage at -80 °C.

### SEC-MALS

Protein samples were characterized using a Superdex75 10/300 Increase column (Cytiva) coupled to a Dawn Heleos II device (Wyatt Technologies) at the joint IBMB/IRB Crystallography Platform, Barcelona Science Park (Catalonia). Analyses were performed at 25ºC, in 20 mM HEPES pH 7.4, 150 mM NaCl buffer at a 0.5 mL/min flow-rate. Bovine serum albumin (BSA) was used as analytical standard. Data were processed and analysed using ASTRA 7 software (Wyatt Technologies).

### Crystallization of Sponge Ig-like proteins

RTK was concentrated to 4.3 µg/µl and SAML to 6.5 µg/µl. Each protein was screened against more than 700 crystallization conditions by the sitting-drop vapor diffusion method. RTK showed the best crystals in 0.2 M ammonium sulfate, 26% w/v PEG4000. The best crystals of SAML were obtained in 0.2 M ammonium sulfate, 30% w/v PEG4000. Crystals were cryoprotected with increasing concentrations of glycerol or ethylene glycol up to 20% in the mother liquor before being cryo-cooled in liquid nitrogen prior to diffraction data analysis.

### Data collection and refinement

Crystals were diffracted at the BL13-XALOC beamline of the ALBA Synchrotron (Cerdanyola del Vallès, Barcelona, Spain). Collected datasets of RTK and SAML were processed with autoPROC^41^ and merged and scaled with *AIMLESS*^42^. SAML was solved using molecular replacement with Phaser MR^43^, using as templates individual Ig-domains from the Alphafold predictions (AF-Q9U965-F1), which were obtained from Uniprot. The RTK structure was obtained via molecular replacement using as template the previously solved SAML structure. A restrained refinement was carried out using phenix.refine^44^ or refmac5^45^. The final atomic coordinates were obtained after successive rounds of refinement and manual building in Coot^46^.

## Supporting information

Full list tandemers

## Acknowledgements

We are grateful to Gilda Dichiara Rodríguez and Sergio Morales Hernández for their technical support. We also thank the staff of the Xaloc beamline at ALBA Synchrotron for their assistance with X-ray diffraction data collection.

## Author contributions

Conceived research project: JLS; Performed experiments: AU, JLS, PY; Data analysis: AU, JLS, JW, JS, PY, EE; Draft writing: JLS, PY.

## Data availability

Atomic coordinates and structure factors for SAML and RTK have been deposited in the Protein Data Bank under the accession codes 8OVQ and 8QPX, respectively.

