## Supplementary figures and images for "Unusual traits shape the architecture of the Ig ancestor molecule"

### Supplementary Figure 1

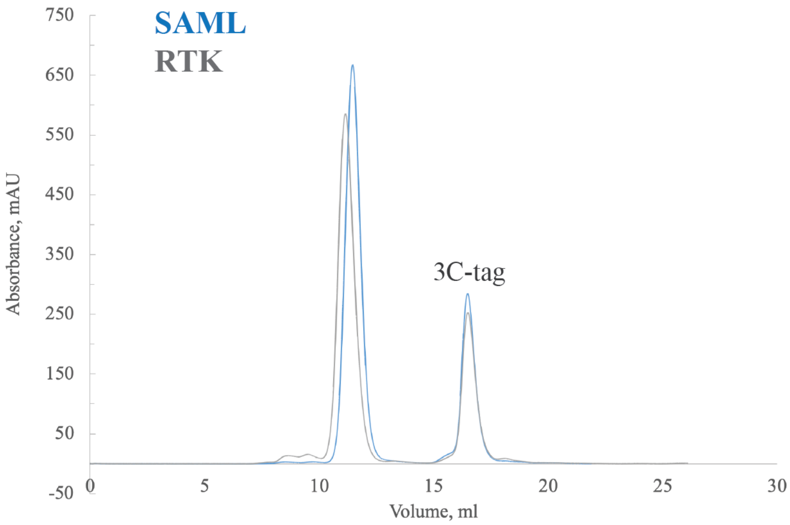

### Supplementary Figure 2

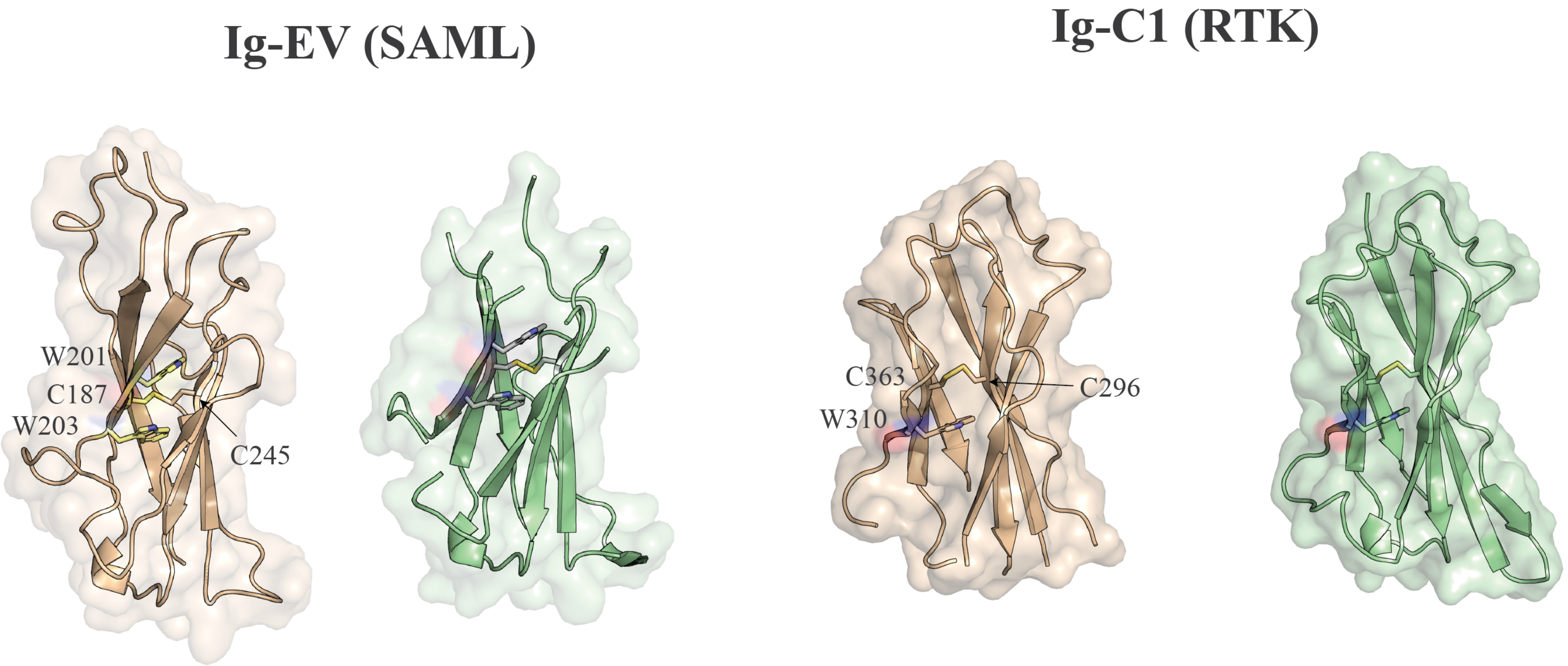

### Supplementary Figure 3

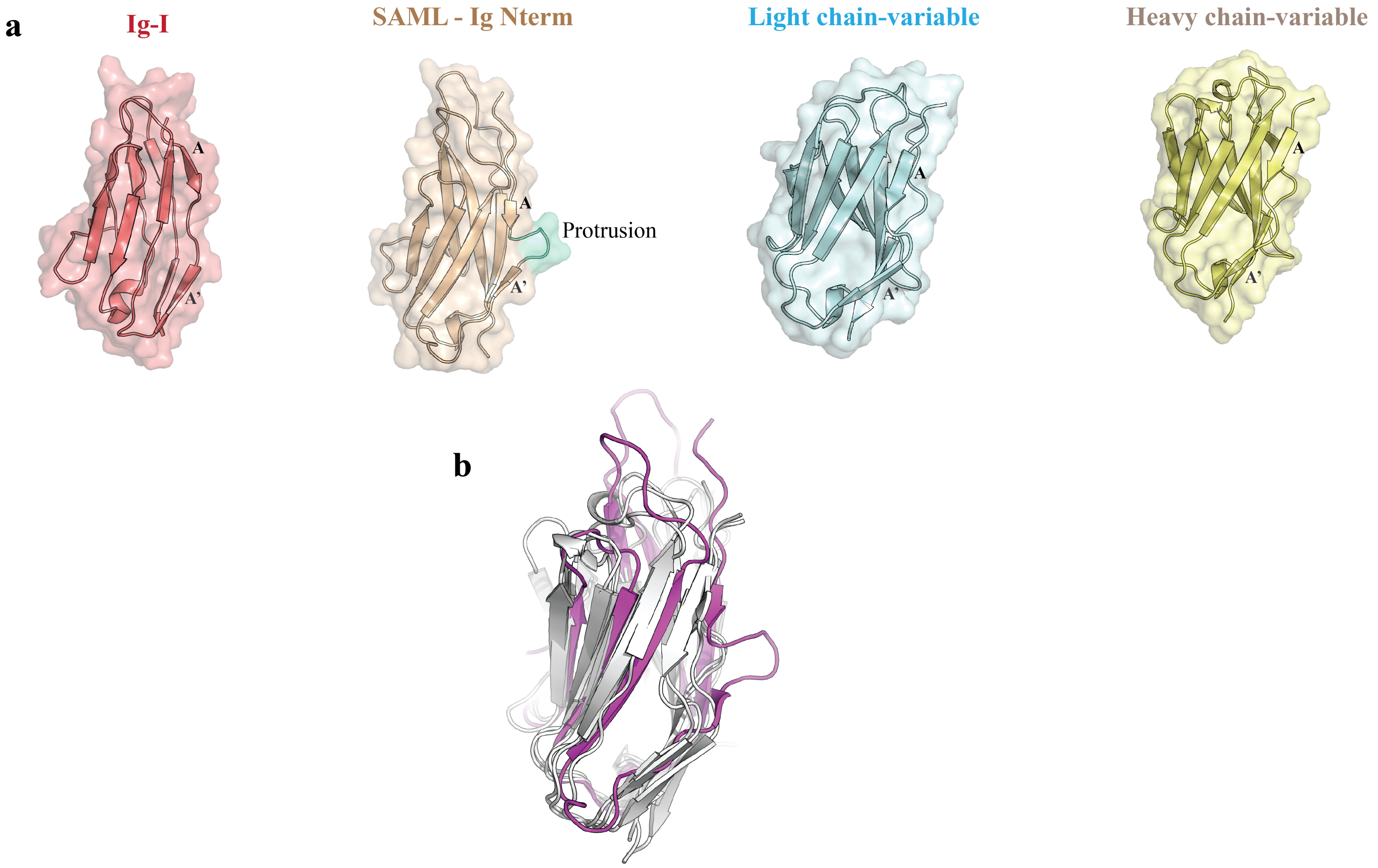

### Supplementary Figure 4

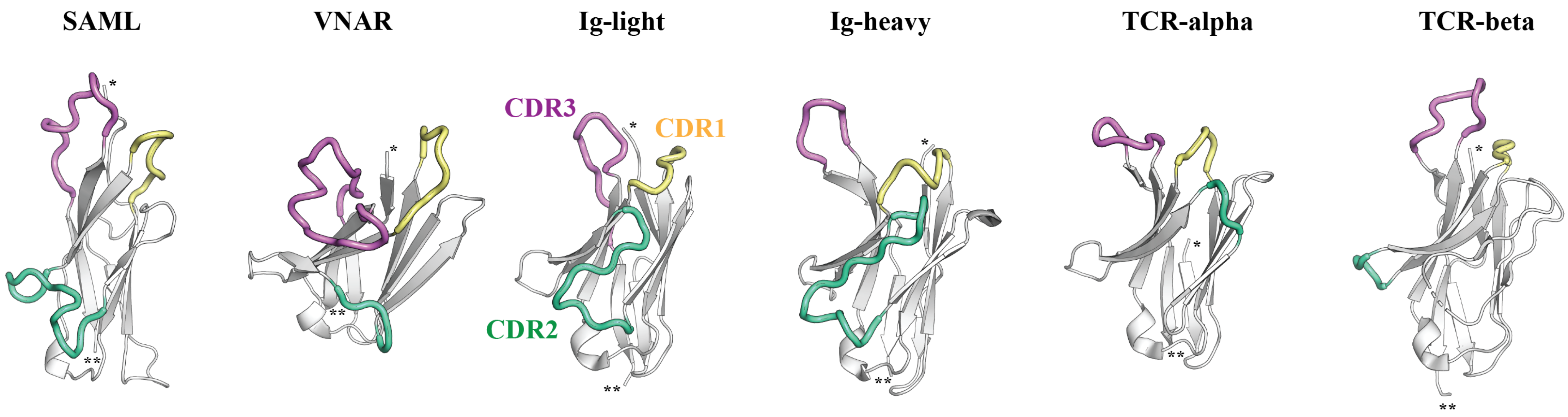

### Supplementary Figure 5

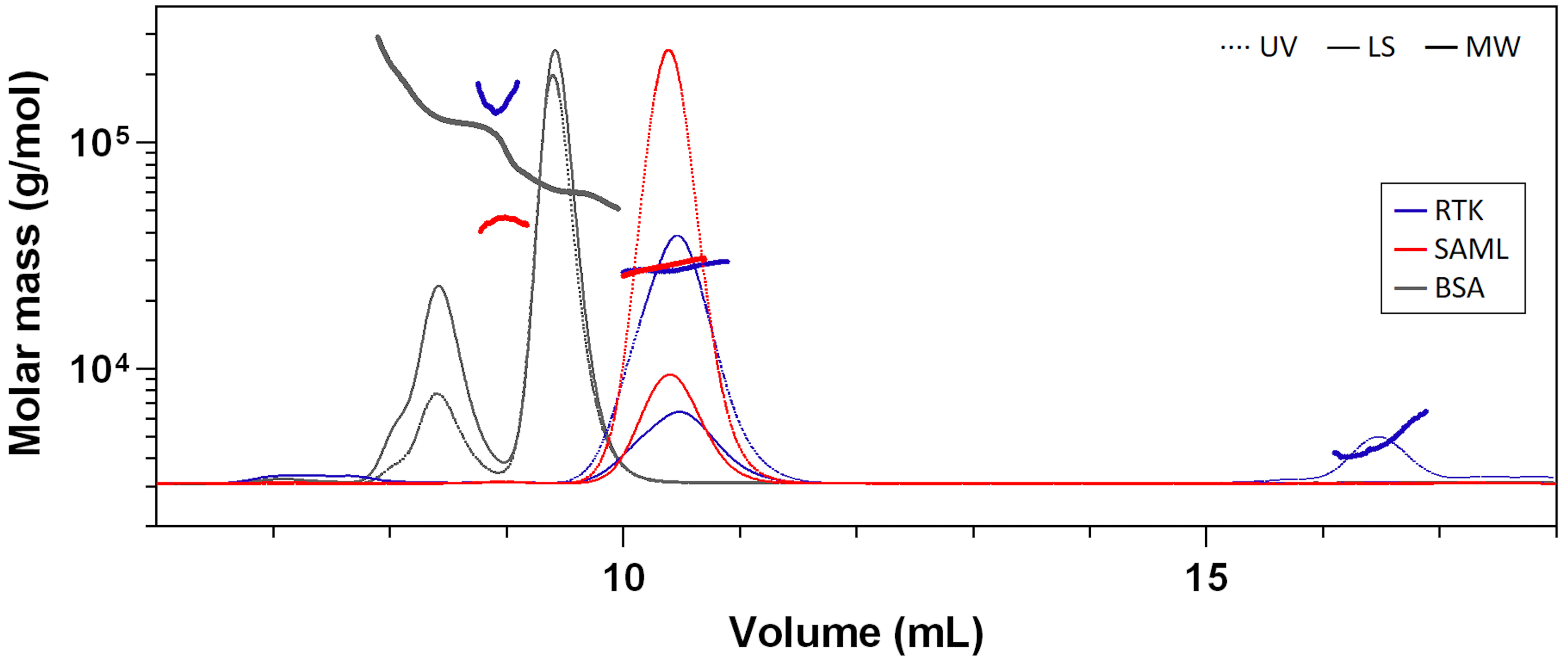

### Supplementary Figure 6

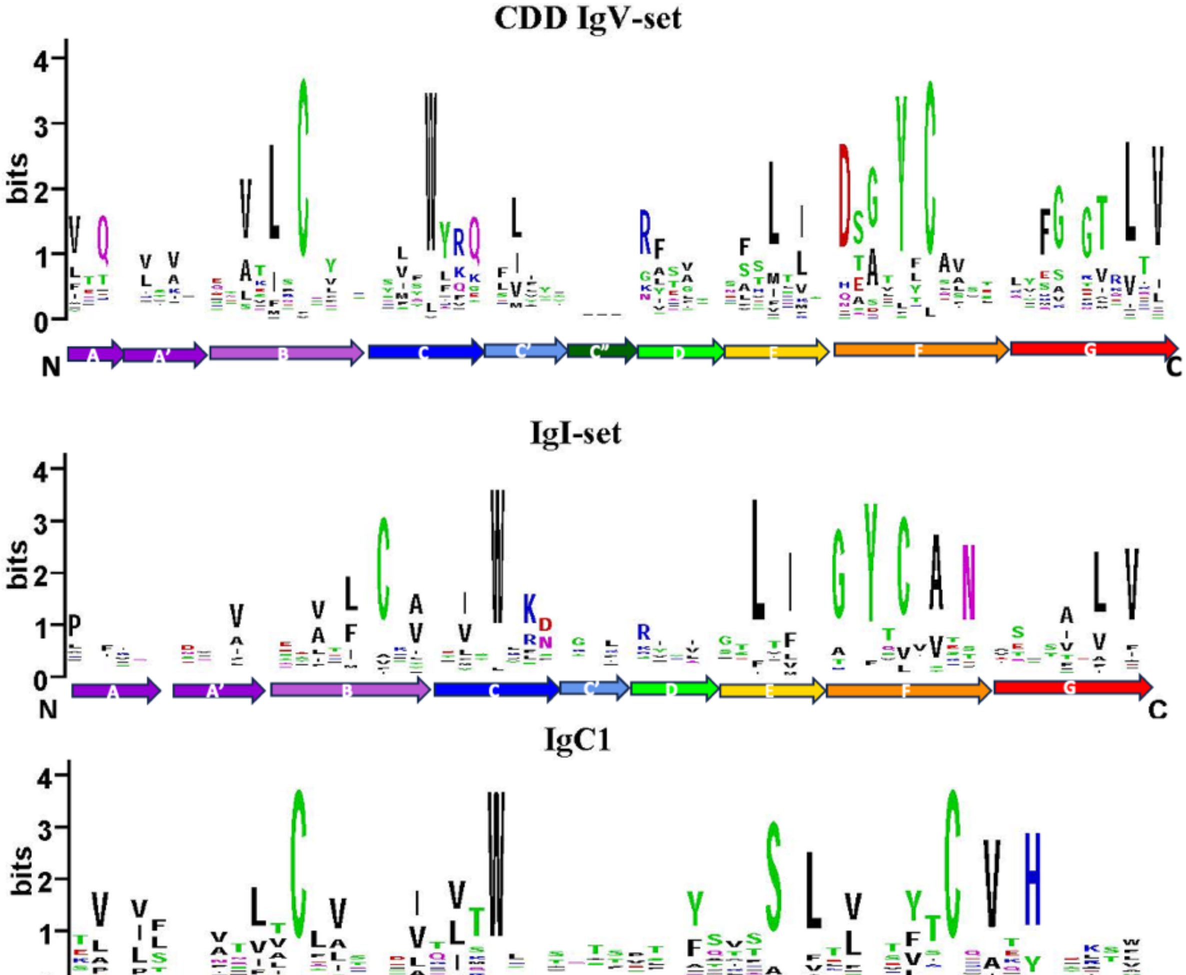

### Supplementary Figure 7

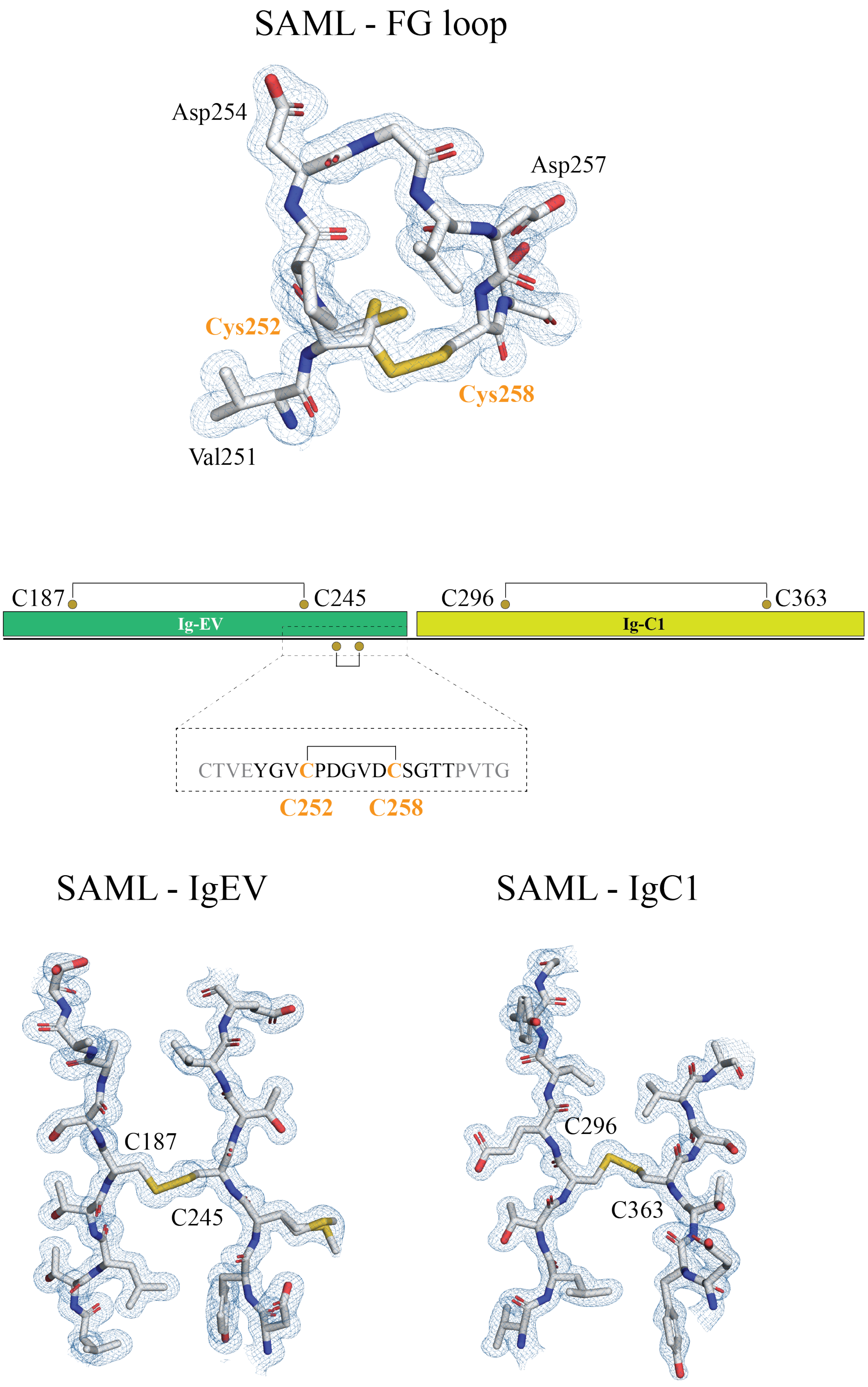

### Supplementary Figure 8

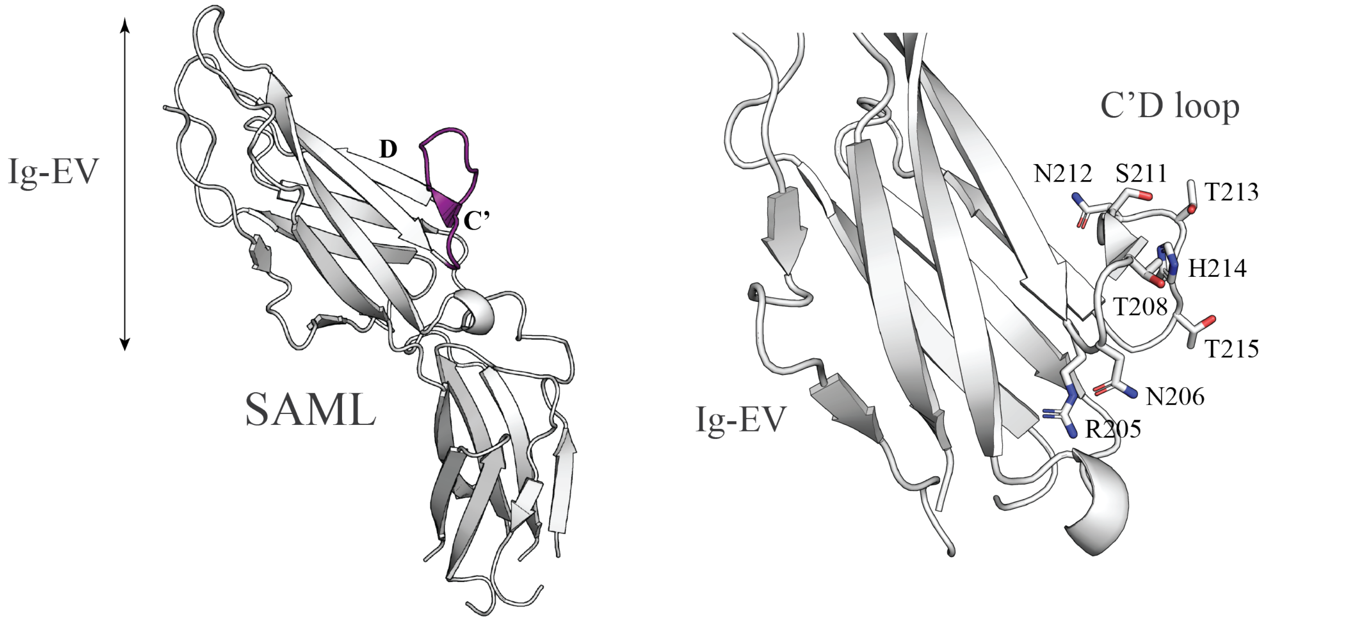


| | | | | | | | | C' strand | | | | C'-D loop | | | | | D strand | | | | | | | | | |
|---|---|---|---|---|---|---|---|---|---|---|---|---|---|---|---|---|---|---|---|---|---|---|---|---|---|---|
| **SAML** | Q | W | Q | W | R | R | N | G | T | L | L | S | N | T | H | T | R | F | S | I | T | P | S | T | N | T |
| **RTK** | Q | W | Q | W | R | R | N | G | T | L | L | S | N | T | H | T | R | F | S | I | T | P | S | T | N | T |
| | . | . | . | . | . | . | . | . | . | . | . | . | . | . | . | . | . | . | . | . | . | . | . | . | . | . |
| | 200 | | | | | 205 | | | | | 210 | | | | | 215 | | | | | 220 | | | | | 225 |

### Supplementary Figure 9

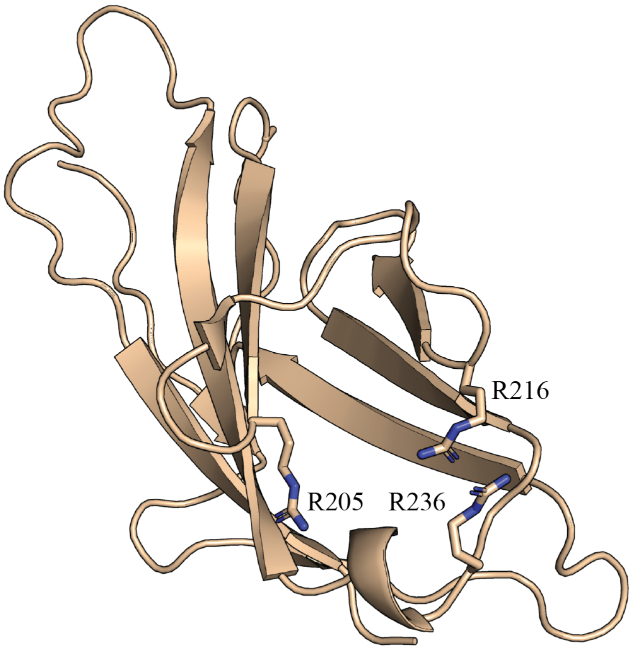

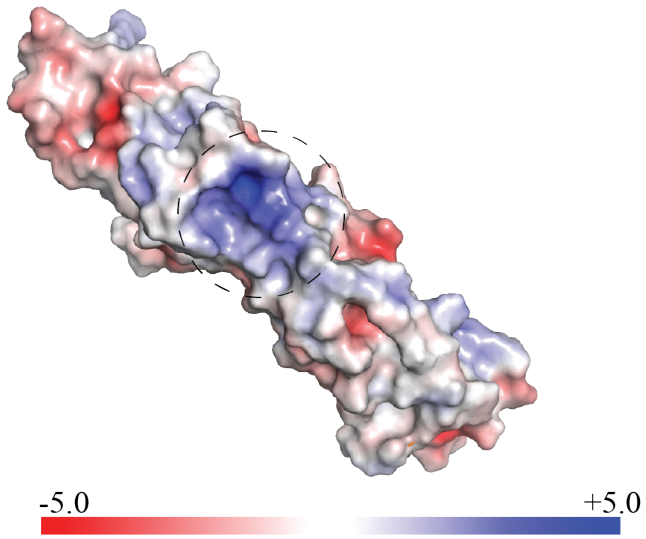

### Supplementary Figure 10

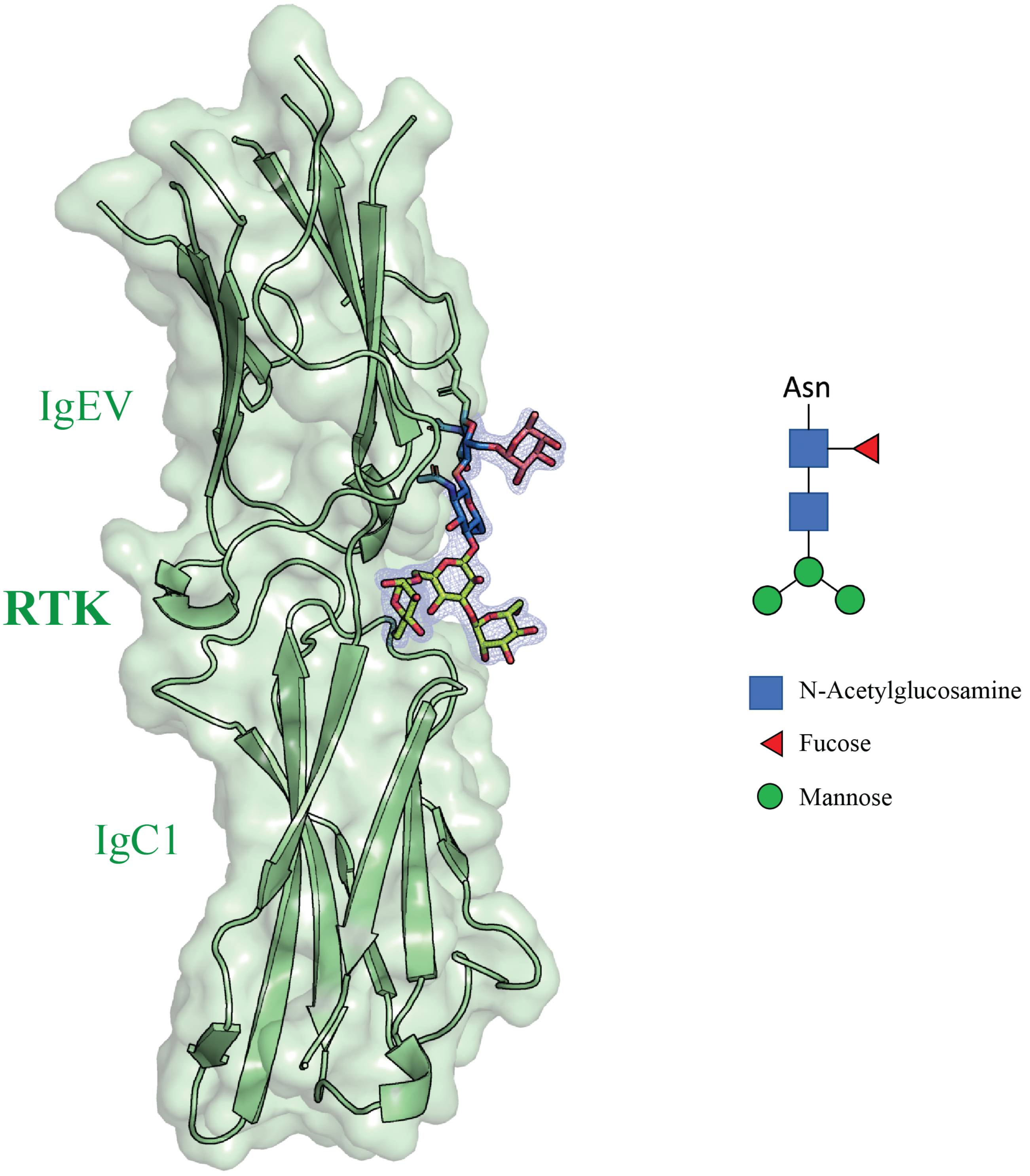
