## Supplementary Material for "Unusual traits shape the architecture of the Ig ancestor molecule"

[1] Unit of Protein Crystallography and Structural Immunology, Navarrabiomed, 31008, Navarra, Spain.

[2] Public University of Navarra (UPNA), Pamplona, 31008, Navarra, Spain.

[3] Navarra University Hospital, Pamplona, 31008, Navarra, Spain.

[4] National Center for Biotechnology Information, National Library of Medicine, National Institutes of Health, Bethesda, MD, United States.

[5] Cancer Data Science Laboratory, Center for Cancer Research, National Cancer Institute, National Institutes of Health, Bethesda, MD, United States.

[*] Equal contribution.

[‡] To whom correspondence should be addressed:

.

Supplementary Figures 1 to 10

Supplementary Table 1

**Supplementary Figure 1. SAML and RTK elution profiles under size exclusion chromatography.** Both SAML and RTK were purified via their N-terminal TwinStrep tag. The proteins were treated with 3C protease to remove the tag and the digestion product was loaded onto a Superdex™ 75 Increase 10/300 GL column. Both proteins eluted as single and symmetric peaks with coherent retention times based on their respective molecular weights. A second peak with larger retention time corresponds to the 3C-released TwinStrep tag.

**Supplementary Figure 2. Conserved disulfide bridges in the Ig b sandwhich.** The N-term Ig-EV and C-term Ig-C1 domains of SAML (wheat color) and RTK (palegreen color) are displayed in cartoon and surface modes. The structurally-conserved cysteines that participate in fundamental disulfide bridges in the Ig domains are highlighted as sticks. Trp residues surrounding them are also highlighted in a similar manner, with W203 and W310 being highly conserved across the Ig superfamily.

**Supplementary Figure 3. Bulge or protrusion observed in SAML/RTK N-terminal Ig domain.** The structures of an Ig-I domain (PDB 2DM3), SAML (PDB 8OVQ and the variable regions of both a human antibody light and heavy chains (PDB 5O1416) were structurally aligned. **a**, shown are the cartoon and surface representations of these molecules preserving in all cases the same orientation with respect to SAML. **b**, overall alignment and of all molecules in grey color and SAML in magenta.

**Supplementary Figure 4. Structural positioning of the CDR regions in primitive and current Ig-domains.** The CDRs present in antibody light and heavy chains as well as in the TCR alpha and beta chains are highlighted in colors. The equivalent regions in the less evolved forms found in *Porifera* SAML and the shark VNAR proteins are also shown for comparison purposes. $^{*}$N_term, $^{**}$C_term.

**Supplementary Figure 5. SEC-MALS analysis of SAML and RTK extracellular regions.** The thick lines represent the molar mass measured by MALS, the dashed lines correspond to SEC-UV peaks, while the solid lines show the Rayleigh ratio. The chromatogram exhibits high homogeneity of the separated monomer fractions of each protein.

**Supplementary Figure 6.** Sequence logos of IgV, IgI and IgC1 domains illustrate the degree of conservation in various strands, as defined by the CDD IgSF. On the y-axis, the probability score where a score of 4 indicates 100% conservation. The x-axis highlights the amino acid positions in strands, using the standard nomenclature A/A',B,C,C',C",D,E,F. IgV domains.

**Supplementary Figure 7. Canonical and non-canonical disulfide bridges in the ancestral SAML and RTK Ig molecules.** All residues displayed in this figure belong to SAML structure and are shown as sticks with carbons, nitrogens, oxygens and sulfurs colored grey, blue, red and yellow, respectively. The 2Fo-Fc electron density maps are shown as blue meshes at a 1 $\sigma$ contour level. A sequence diagram for the extracellular region of SAML is located in the middle panel, and highlights the position of the non-canonical cysteines (yellow circles) in the FG loop (CDR3) of the IgEV domain (upper panel). Their canonical counterparts in the IgEV and IgC1 domains are also shown in the bottom panel. The numbering follows that of the native full-length protein (Uniprot Q9U965).

**Supplementary Figure 8. The C'D loop conformation.** SAML and RTK present a remarkable C'D loop, shown here in violet color in SAML overall structure (left), and in a zoomed in view at right. Underneath is an amino acid alignment of the C'D loop in both SAML and RTK, as well as surrounding residues.

**Supplementary Figure 9. Arginine-rich region in the Ig-EV domain of SAML and RTK.** An arginine-rich area is conserved in SAML and RTK and features a presumably positively-charged and functional pocket. Left, cartoon representation with the three conserved arginines highlighted as sticks. Right, electrostatic surface map of both Ig domains in SAML structure. The dashed circle denotes the positively charge area.

**Supplementary Figure 10. N-glycosylation in Ig-EV.** A site of glycosylation is present in the Ig-EV domains of SAML and RTK. Shown is the cartoon and surface representation of RTK structure with the N-glycosylation patterns displayed as stick. The 2Fo-Fc electron density map, contoured at 1.0 $\sigma$, is indicated as a light blue mesh for the sugar moiety. A glycosylation pattern diagram compatible with N-glycosylation in insect cells as well as the color legend of the sugar molecules are indicated.

**Supplementary Table 1. Selected tandemer homologs.** The first set of homologs (blue) correspond to high ranking tandemer hits in the non-Ab dataset with an IgV-IgC1 topology, followed in grey by a VNAR (for which only single domain structures for the V and C1 domains are available), the CD4 hit correspond to an IgV+IgC2 topology that form a compact tandemer. The second set of homologs (pink) corresponds to high ranking tandemer hits in the Ab/TCR dataset wich all have a V+C1 topology. Finally, the last set of homologs (blue) shows high ranking hits with a diversity of Ig-Ig tandemers containing IgI, IgC2 and unspecified Ig-like topologies. (PdbID = PDB accession; chain = chain identifier; D1 = domain number on PDB structure; Naligned1 = number of aligned residues for domain 1; Cover1 = coverage of SAML domain 1 (Naligned1/number of residues in SAML domain 1 with coordinates); RMSD1 = Rmsd for domain 1; Nid1 = number of identical residues in alignment on domain 1; analogously for D2, Naligned2, Cover2, RMSD2, Nid2).

| PDBid | chain | D1 | Naligned1 | Cover1 | RMSD1 | Nid2 | D2 | Naligned2 | Cover2 | RMSD2 | NId 2 | Protein name | %id1 | %id2 | %id | Name | Topology |
|---|---|---|---|---|---|---|---|---|---|---|---|---|---|---|---|---|---|
| 6EG0 | A | 1 | 80 | 0.69 | 1.931 | 18 | 2 | 62 | 0.674 | 1.933 | 20 | Defective_proboscis_extension_response_4 | 23% | 32% | **27%** | **DPR4** | V+C1 |
| 5ZO2 | A | 1 | 62 | 0.534 | 1.629 | 12 | 2 | 61 | 0.663 | 1.214 | 17 | Cell_adhesion_molecule_4 | 19% | 28% | **24%** | **necl-4** | V+C1 |
| 8DGG | A | 1 | 82 | 0.707 | 2.547 | 18 | 2 | 66 | 0.717 | 1.948 | 10 | Lymphocyte_activation_gene_3_protein | 22% | 15% | **19%** | **LAG-3** | V+C1 |
| 4BFI | B | 1 | 82 | 0.707 | 1.965 | 18 | 2 | 60 | 0.652 | 2.213 | 11 | OX-2_MEMBRANE_GLYCOPROTEIN | 22% | 18% | **20%** | **CD200** | V+C1 |
| 3EPD | R | 1 | 95 | 0.819 | 2.661 | 21 | 2 | 89 | 0.967 | 4.946 | 13 | Poliovirus_receptor | 22% | 15% | **18%** | **CD155** | V+C1 |
| 1I8L | B | 1 | 72 | 0.621 | 2.151 | 19 | 2 | 87 | 0.946 | 3.421 | 8 | T_LYMPHOCYTE_ACTIVATION_ANTIGEN_CD80 | 26% | 9% | **18%** | **CD80** | V+C1 |
| 3BIK | A | 1 | 76 | 0.655 | 1.825 | 13 | 2 | 70 | 0.761 | 2.534 | 11 | Programmed_cell_death_1_ligand_1 | 17% | 16% | **16%** | **PDL1** | V+C1 |
| 3BP5 | B | 1 | 76 | 0.655 | 1.987 | 10 | 2 | 61 | 0.663 | 2.007 | 10 | Programmed_cell_death_1_ligand_2 | 13% | 16% | **15%** | **PDL2** | V+C1 |
| 4FRWB | B | 1 | 75 | 0.647 | 2.002 | 15 | 1 | 66 | 0.717 | 2.216 | 6 | Poliovirus_receptor-related_protein_4 | 20% | 9% | **15%** | **nectin-4** | V+C1 |
| 6YJP | C | 1 | 73 | 0.629 | 2.538 | 13 | 2 | 61 | 0.663 | 1.854 | 12 | Natural_cytotoxicity_triggering_receptor_3_ligand_1 | 18% | 20% | **19%** | **B7-H6** | V+C1 |
| 1VES | A | 1 | 93 | | 3.39 | | | | | | | Antigen Receptor variable domain from sharks | 17% | | | VNAR | V+C1 |
| 1WIQ | B | 1 | 77 | 0.664 | 1.901 | 19 | 2 | 74 | 0.804 | 2.753 | 9 | T-CELL_SURFACE_GLYCOPROTEIN_CD4 | 25% | 12% | **18%** | **CD4** | V+C2 |
| 1SEQ | L | 1 | 74 | 0.638 | 1.809 | 29 | 2 | 65 | 0.707 | 2.086 | 12 | Monoclonal_Antibody_MNAC13 | 39% | 18% | **29%** | **VL** | V+C1 AB L |
| 8BBH | H | 1 | 80 | 0.69 | 2.211 | 16 | 2 | 59 | 0.641 | 1.851 | 19 | Heavy_Chain_of_TL1_Fab_fragment | 20% | 32% | **26%** | **CH1** | V+C1 AB H |
| 6ULRE | E | 1 | 74 | 0.638 | 1.852 | 20 | 2 | 60 | 0.652 | 1.521 | 13 | TCR-V-beta_5-6*01 | 27% | 22% | **24%** | **TCRb** | V+C1 TCRb |
| 7XY8 | A | 1 | 64 | 0.552 | 2.853 | 14 | 2 | 72 | 0.783 | 2.61 | 21 | Isoform_2_of_Basigin | 22% | 29% | **26%** | **CD147** | |
| 5VKJ | A | 2 | 78 | 0.672 | 2.5 | 20 | 3 | 50 | 0.543 | 1.141 | 12 | B-cell_receptor_CD22 | 26% | 24% | **25%** | **CD22** | |
| 5JDD | A | 1 | 78 | 0.672 | 1.732 | 12 | 2 | 62 | 0.674 | 2.413 | 19 | Titin | 15% | 31% | **23%** | **Titin** | |
| 2X1X | R | 1 | 70 | 0.603 | 2.891 | 9 | 2 | 66 | 0.717 | 1.659 | 20 | VASCULAR_ENDOTHELIAL_GROWTH_FACTOR_RECEPTOR_2 | 13% | 30% | **22%** | **VEGF2** | |
| 8DPU | L | 1 | 49 | 0.422 | 1.177 | 16 | 2 | 57 | 0.62 | 1.994 | 5 | Interleukin-11_receptor_subunit_alpha | 33% | 9% | **21%** | **IL11Ra** | |
| 5OJ2 | A | 3 | 69 | 0.595 | 2.208 | 16 | 4 | 63 | 0.685 | 1.229 | 11 | MAM_domain-containing_glycosylphosphatidylinositol_anchor_protein_1 | 23% | 17% | **20%** | **MDGA1** | |
| 6UEA | C | 2 | 93 | 0.802 | 3.1 | 21 | 3 | 63 | 0.685 | 2.554 | 7 | Polymeric_immunoglobulin_receptor | 23% | 11% | **17%** | **PIgR** | |
| 4OFY | E | 2 | 97 | 0.836 | 3.496 | 15 | 3 | 58 | 0.63 | 1.655 | 8 | Protein_SYG-2 | 15% | 14% | **15%** | **SYG-2** | |
| 7R5K | 15 | 1 | 69 | 0.595 | 2.967 | 11 | 2 | 46 | 0.5 | 1.148 | 5 | Nuclear_pore_membrane_glycoprotein_210 | 16% | 11% | **13%** | **Pom210** | |

**Supplementary Table 2. Full list tandemer homologs.** Full hits set of 15,000 2-domain (or more) chains (first two domains) with max %id ca. 0.47 on Ig 1 (VL) (based on structurally aligned residues) and max %id ca. 0.33 on Ig 2.
